# MUC4-targeted CAR-T Cells Restrain Chemotherapy-resistant Colorectal Cancer

**DOI:** 10.64898/2026.02.26.708339

**Authors:** Avik Chattopadhyay, Erin C. O’Connor, Anna M. Tingler, Kristi L. Helke, Melinda A. Engevik, Leonardo M.R. Ferreira

## Abstract

Colorectal cancer (CRC) is the leading cause of cancer mortality in young adults, with chemoresistance and metastatic progression severely limiting treatment options. MUC4, a membrane-bound mucin normally confined to the apical surface of epithelial cells, becomes aberrantly overexpressed across the tumor cell membrane in CRC and other cancer types, promoting cancer survival, immune evasion, and metastasis. Therefore, MUC4 is a promising candidate for the development of targeted therapies. We detected MUC4 cell surface expression in a panel of human CRC cell lines. We engineered MUC4-specific chimeric antigen receptor (CAR) T cells and evaluated their activity against methotrexate-resistant MUC4-expressing human CRC cell lines HT29-MTX and T84. MUC4 CAR-T cells efficiently eliminated HT29-MTX and T84 cells *in vitro* and delayed HT29-MTX subcutaneous tumor growth *in vivo*. Importantly, MUC4 CAR-T cell treatment significantly reduced tumor burden and improved survival in a highly aggressive and lethal CRC model, intraperitoneal HT29-MTX CRC metastatic dissemination in NSG mice. Mouse necropsies revealed no off-target *in vivo* toxicities associated with MUC4 CAR-T cell therapy. Altogether, our findings establish MUC4 as an important and clinically relevant CAR-T cell target and demonstrate therapeutic efficacy of MUC4 CAR-T cell therapy for invasive CRC, a setting where chemotherapy typically fails. MUC4-targeted CAR-T cell therapy has the potential to fill a critical treatment gap for patients with chemoresistant and invasive CRC and may extend to other MUC4-expressing solid tumors.

**Statement of Priority:** Colorectal cancer is increasingly diagnosed in younger individuals, a trend accompanied by a higher prevalence of aggressive and chemotherapy-resistant disease. Despite advances in systemic therapy, outcomes for patients with refractory or metastatic colorectal cancer remain poor, underscoring a critical unmet clinical need. CAR-T cell therapy offers a mechanism to selectively target tumor cells that evade conventional treatments and may thus be particularly valuable in mucinous, chemotherapy-resistant subsets. Developing effective CAR-T strategies for new targets in colorectal carcinoma, such as MUC4, is an urgent priority in this era of rising early-onset disease.

## Introduction

Colorectal cancer (CRC) incidence has risen sharply among adults aged 45-49 years, with annual increases accelerating from 1.1% during 2004-2019 to 12.0% during 2019-2022. These numbers have been driven primarily by local-stage tumors. Advanced-stage disease has risen steadily in individuals younger than 45 and in those aged 45–54 years, underscoring the urgent need for novel therapeutic strategies (1). Within CRC cases, mucinous CRCs are frequently associated with metastatic dissemination, poor prognosis, and resistance to conventional chemotherapies. Despite advances in targeted agents and immune checkpoint blockade, durable responses in metastatic CRC, especially in chemotherapy-refractory disease, remain limited (2).

Mucins are high–molecular–weight, heavily glycosylated secreted and transmembrane proteins that contribute to epithelial protection but are frequently dysregulated in malignancies (3). In CRC and other solid tumors, mucins become aberrantly expressed with the loss of cell polarity and gain of tumor-aggressive properties and promote tumor progression, immune evasion, and therapeutic resistance (3). Among membrane-bound mucins, MUC1 and MUC16 have been extensively explored as immunotherapy targets; however, their broad baseline expression across normal epithelia and in the hematopoietic compartment raises concerns regarding on-target/off-tumor toxicity (4). In contrast, MUC4 exhibits several features that make it an attractive tumor-associated antigen in several solid tumor types. MUC4 is often markedly upregulated and depolarized in malignant epithelium while showing more restricted expression in many normal tissues, potentially offering an improved therapeutic window (5). Moreover, MUC4 gene polymorphisms and expression levels are associated with CRC prevalence and survival (6,7).

Chimeric antigen receptor (CAR) T cell therapy has revolutionized the treatment of hematologic malignancies by enabling antigen-specific redirection of T cell cytotoxicity, boasting seven FDA approvals and remission rates of up to 90% (8). However, translating CAR-T efficacy to solid tumors has been hindered by target antigen heterogeneity, on-target/off-tumor toxicity, poor tumor trafficking, and the immunosuppressive tumor microenvironment (9,10). Targeting MUC4 in CRC represents a promising strategy to overcome some of these barriers. Here, we developed and functionally characterized second-generation MUC4-targeted CAR-T cells and evaluated their efficacy against chemotherapy-resistant human CRC models *in vitro* and *in vivo*.

## Methods

### Molecular biology and lentivirus production

MUC4CAR-2A-NGFRt and Luciferase-2A-GFP lentiviral plasmids were synthesized by VectorBuilder (Chicago, IL). All genes were driven by an EF1A promoter. The MUC4CAR gene contained a CD8a signal peptide, an N-terminal Myc-tag, an scFv sequence derived from antibody clone 8G7 recognizing human MUC4, a CD8 hinge domain, a CD28 transmembrane domain, and a CD28-CD3z signaling domain (11). Lentivirus was produced by transfecting HEK293FT cells with the plasmid of interest and lentiviral packaging plasmids, as previously described (12). Lentivirus was titrated, aliquoted, and stored at -80 °C until use.

### Cell culture

HT-29-MTX, T84, and HEK293T cells were used between passages 50 and 100 for the first two cell lines and up to passage 25 for HEK293FT cells. All cell lines were routinely grown in DMEM-10, composed of DMEM (25 mM glucose) basal media supplemented with 10% heat inactivated (30 min. 56°C) fetal bovine serum (FBS) and penicillin-streptomycin (all from Gibco, ThermoFisher Scientific, Waltham, MA). Cell lines were passaged using 0.25% Trypsin-EDTA (Gibco), seeded at 1 x 10^6^ cells per 75 cm^2^ flasks, and kept in a tissue culture incubator at 37 °C in a 5% CO_2_, 95% air atmosphere. For maintenance purposes, cells were passaged every 2–3 days to maintain logarithmic growth. All experiments were performed in 24-well (5 × 10□ cells per well) or 96-well plates (1 × 10□ cells per well). Primary human T cells were cultured in RPMI-10, composed of RPMI-1640 basal media supplemented with 10% heat inactivated FBS, GlutaMAX, sodium pyruvate, non-essential amino acids (NEAA), HEPES, and penicillin-streptomycin (all from Gibco) in the presence of 100 IU/ml recombinant human IL-2 (PeproTech, ThermoFisher Scientific). Cells were passaged and fresh medium and IL-2 added every other day.

### CAR-T cell generation

Human T cells were isolated from peripheral blood mononuclear cells (PBMCs) obtained from de-identified blood donors (STEMCELL Technologies, Vancouver, BC, Canada) by magnetic negative selection using a Human CD3^+^ T Cell Isolation Kit (STEMCELL Technologies) following manufacturer’s instructions and activated with anti-CD3/CD28 dynabeads (Gibco) at a 1:1 bead to cell ratio and 100 IU/ml IL-2. Two days after activation, T cells were transduced with MUC4CAR lentivirus at a multiplicity of infection of 1 (one particle per cell) in the presence of 100 IU/ml IL-2. Immediately after adding the lentivirus, T cells were centrifuged at 1,000 x g at 32 °C for 1 h and then returned to the tissue culture incubator, as previously described (12). Following transduction, T cells were maintained and expanded in RPMI-10 medium with fresh medium and IL-2 being given every 2 days. CAR-expressing T cells were purified by magnetic positive selection based on reporter NGFRt expression using a Human CD271 Positive Selection Kit (STEMCELL Technologies), as per manufacturer’s instructions, before being used in experiments.

### Flow cytometry

Cell lines and T cells were surface stained with antibodies in phosphate buffered saline (PBS) (Gibco) in FACS tubes for 30 min at 4°C in the dark, washed, and resuspended in PBS for reading in a CytoFLEX flow cytometer (Beckman Coulter, Brea, CA). Antibodies used in this study were MUC4 (clone 87G, Santa Cruz Biotechnology, Santa Cruz, CA), goat anti-mouse AlexaFluor647 (ThermoFisher), Myc-tag AlexaFluor647 (Cell Signaling Technology, Danvers, MA), and NGFR/CD271 PE (Biolegend, San Diego, CA), as well as Ghost Red780 viability dye (Cytek, Fremont, CA). Flow cytometry data were analyzed using FlowJo v10.0 (FlowJo, Ashland, OR).

### Luciferase-labeled cancer cell generation

Lentiviral transduction of the cell lines with the dual reporter construct encoding firefly luciferase and GFP separated by a 2A sequence was performed at different multiplicities of infection (MOIs). Among the conditions tested, an MOI of 10 yielded optimal and uniform reporter gene integration without compromising cell viability. Following transduction, GFP-positive cells were sorted by fluorescence-activated cell sorting (FACS) using a BD FACS Aria sorter (Becton Dickinson, Franklin Lakes, NJ) and subsequently expanded over multiple passages. FACS-sorted T84-Luc-GFP and HT29-MTX-Luc-GFP cells maintained stable GFP expression, suggesting durable lentiviral integration and transcriptional stability. To confirm functional luciferase activity, the engineered cells were incubated with D-luciferin substrate (Biosynth, Gardner, MA) and imaged for photon emission using an AMI bioluminescence detection system (Spectral Instruments Imaging, Tucson, AZ). Both T84-Luc-GFP and HT29-MTX-Luc-GFP cell lines emitted robust and quantifiable bioluminescence signals, validating their suitability for *in vivo* tracking.

### Cytotoxicity measurement

Target cancer cells were seeded at a density of 10^4^ cells/well in a 96-well flat-bottom plate and co-cultured with MUC4CAR-T or UT-T cells at different effector-to-target (E:T) ratios for 36 hours. At the end of the co-incubation, three different techniques were employed to measure cytoxicity. CyQUANT Lactate dehydrogenase (LDH) release cytotoxicity assay was performed according to the manufacturer’s instructions (ThermoFisher). At the end of incubation, 50 μL of culture supernatant were carefully harvested and transferred to a new plate, after which LDH detection reagent was added. After color development, absorbance at 490 nm was measured using a microplate reader (SpectraMax, Molecular Devices, San Jose, CA). Percent cytotoxicity was calculated relative to spontaneous and maximum lysis controls, allowing normalization across experiments. All conditions were performed in technical triplicates. CCK-8 viability (cytotoxicity) assay was performed according to the manufacturer’s instructions (Millipore-Sigma, Burlington, MA). At the end of incubation, media was aspirated, adherent cancer cells were washed with PBS, and fresh media was added with 10 μL of CCK-8 reagent was added directly to each well with a total volume of 100 μL and incubated at 37 °C to allow WST-8 reduction by metabolically active cells. Absorbance at 450 nm was measured using a microplate reader, and relative viability was calculated after background subtraction and normalization to target-only controls. Reduced metabolic signal was interpreted as increased CAR-T cell-mediated cytotoxicity. Cell viability was also determined by flow cytometry by harvesting the co-cultures with Trypsin-EDTA, staining the cells with Ghost Red780 viability dye, and quantifying percentage of dye positive cells.

### NSG mouse xenograft tumor models

All experiments were performed in 8-12 weeks old immunodeficient NSG mice (Jackson Laboratories, Bar Harbor, ME). Female NSG mice were subcutaneously injected with 5 x 10^6^ HT29-MTX cells. Tumor measurements were performed using a caliper starting at one week post tumor injection, at which time mice were randomized based on tumor size and received 5 x 10^6^ untransduced (UT) or MUC4 CAR T cells intravenously via retro-orbital injection (13). Tumor size was measured over time with calipers. For the metastatic cancer dissemination model, luciferase modified cells were injected intraperitoneally into NSG mice: 5 x 10^6^ T84-luc-GFP in male mice and 5 x 10^6^ HT29-MTX-luc-GFP in female mice. One week later, luciferase activity measurements started. Mice were injected intraperitoneally with 100 μL of 0.22 mm filter-sterilized D-luciferin substrate at 15 mg/ml and, 8 minutes later, imaged in the AMI optical imaging system under isoflurane anesthesia for 1 minute (14). Luciferase activity data analysis was performed using Aura software (Spectral Instruments Imaging). Mice were randomized based on luciferase signal and received 5 x 10^6^ UT or MUC4 CAR T cells intravenously via retro-orbital injection. Tumor size was measured over time with calipers. Tumor burden was measured over time using bioluminescence imaging and at the experimental endpoint via mouse necropsy.

### Statistical analysis

Statistical analyses were performed using GraphPad Prism v10.0 (GraphPad Software, San Diego, CA). Data are presented as mean ± SD unless otherwise indicated. Statistical significance was determined using the tests specified in the corresponding figure legends. A p value < 0.05 was considered statistically significant.

## Results

### MUC4 is a cell surface target in colorectal cancer

To test our hypothesis that MUC4 is a new target for chimeric antigen receptor (CAR) T cell therapy against hard-to-treat colorectal cancer (CRC), we first evaluated MUC4 gene expression in different CRC subtypes and stages. We found that MUC4 expression is higher in mucinous compared to non-mucinous CRC across all stages, from stage i (localized) to stage iv (metastatic) (**Fig. 1A**). Next, we sought to survey MUC4 surface expression in a panel of human CRC cell lines. All lines tested expressed MUC4, as determined by flow cytometry with anti-MUC4 clone 8G7 antibody (**Fig. 1B**). Interestingly, two of the highest MUC4 expressors, HT29-MTX and T84, are known to be highly mucinous and resistant to chemotherapeutic agents, such as methotrexate (MTX) (15-17). Given this observation and the fact that chemotherapy resistant CRC represents the biggest need for new therapies, we focused on HT29-MTX and T84 cells in subsequent experiments.

**Figure 1.**
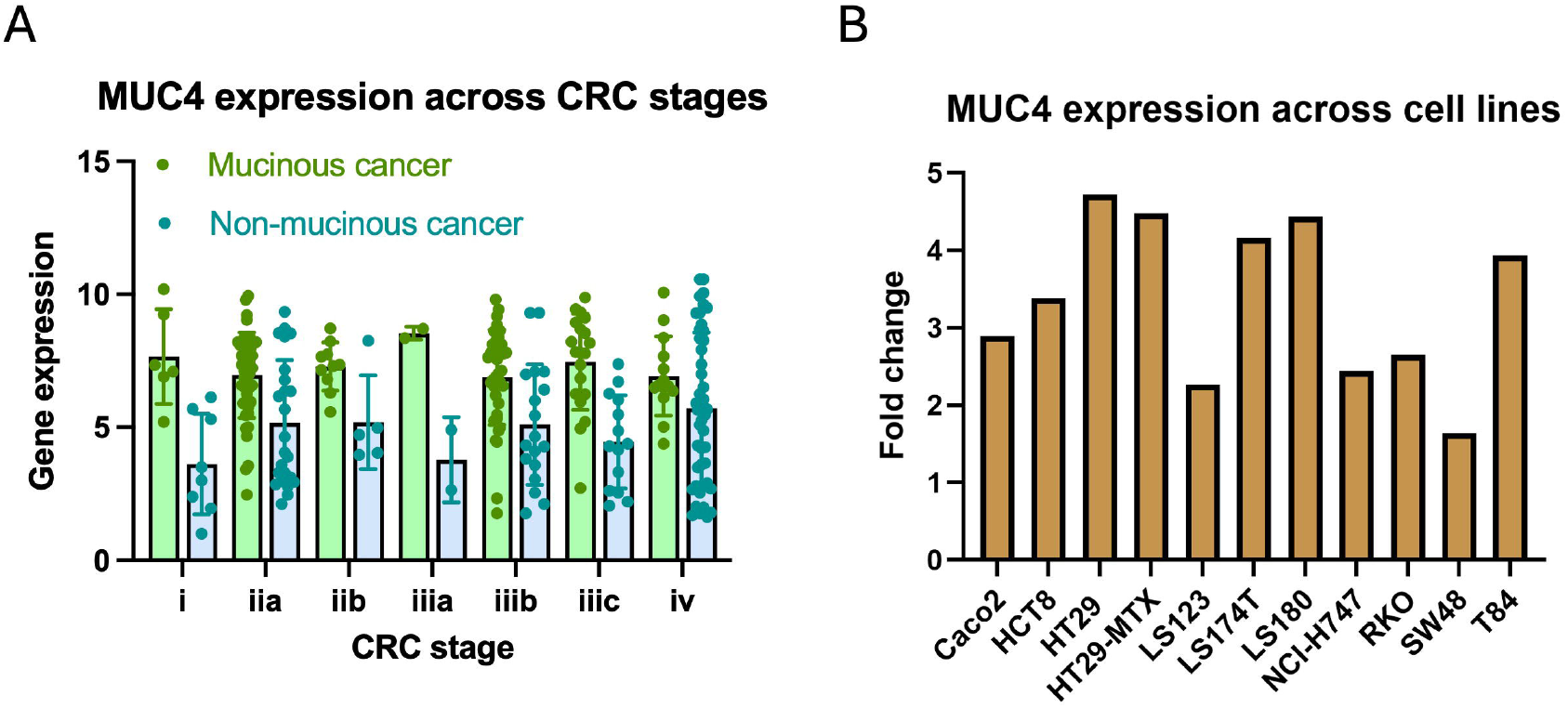
MUC4 is highly expressed in colorectal cancer, especially in chemotherapy-resistant mucinous tumors. (A) MUC4 gene expression in mucinous and non-mucinous colorectal cancer across cancer stages. (B) MUC4 surface expression across colorectal cancer cell lines, as determined by flow cytometry. Fold change of media fluorescence intensity (MFI) of MUC4 staining over IgG1 isotype control.

### MUC4 CAR-T cells mediate potent and selective cytotoxicity against chemotherapy-resistant CRC cells

To target MUC4, we took a single chain variable fragment (scFv) sequence corresponding to anti human MUC4 antibody clone 8G7 and engineered a second-generation CD28-CD3zeta CAR construct targeting MUC4. This construct incorporated a Kozak sequence upstream of a CD8_α_ signal peptide sequence to optimize expression and membrane trafficking. Moreover, an N-terminal Myc-tag was included to facilitate cell surface detection of this new CAR. The antigen-binding domain, comprising the scFv formed by MUC4-specific clone 8G7-derived VL and VH domains linked via a (GGGS)_3_ flexible linker, was fused to a CD8 hinge and a CD28 transmembrane domain, followed by a CD28-CD3zeta tandem intracellular signaling domain. Importantly, a T2A self-cleaving peptide sequence allowed for bicistronic expression of the CAR and a truncated nerve growth factor receptor (NGFRt) reporter gene as an independent and selectable surface marker of T cell transduction (**Fig. 2A**).

**Figure 2.**
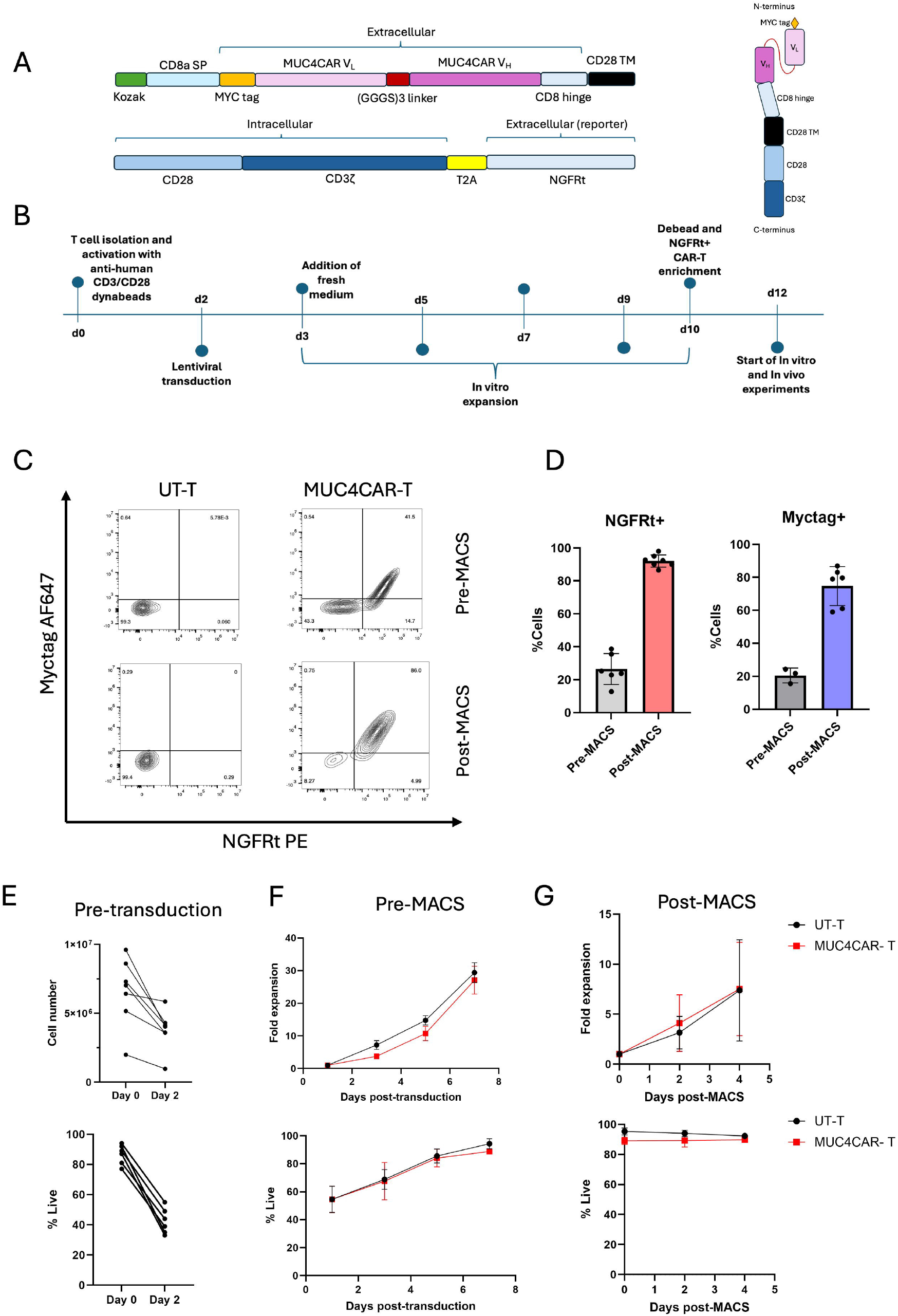
MUC4 CAR-T cells express CAR on the cell surface and retain robust expansion capacity. (A) Schematic representation of second-generation MUC4 CAR DNA construct with T2A-linked NGFRt reporter (left) and MUC4 CAR protein (right). (B) Workflow of primary human T cell activation, lentiviral CAR transduction, magnetic-activated cell sorting (MACS), expansion, and downstream functional assays. (C) Representative flow cytometry plots showing CAR surface (Myc-tag) and reporter (NGFRt) detection at day 6 post-transduction (pre-MACS) and day 2 following MACS enrichment (post-MACS). (D) Quantification of NGFRt^+^ and Myc^+^ T cells after transduction and after MACS enrichment across donors. (E) Total cell number (top) and percent viability (bottom) measured by automated cell counting at day 0 and day 2 following T cell isolation and activation. (F) Expansion kinetics (top) and cell viability (bottom) of MUC4 CAR-T cells following lentiviral transduction (pre-MACS). (G) Expansion (top) and viability (bottom) of MUC4 CAR-T cells after NGFR-based MACS enrichment. Data are presented as mean ± SEM from independent donors.

Primary human T cells were isolated from peripheral blood mononuclear cells (PBMCs), activated with anti-CD3/CD28 beads and 100 IU/ml IL-2, transduced with MUC4 CAR lentivirus, and expanded *in vitro* (**Fig. 2B**). MUC4 CAR was successfully expressed on the surface of T cells, as per Myc-tag staining (**Fig. 2C**) and we achieved initial transduction efficiencies of ∼30% across donors, as assessed by frequency of Myc-tag^+^NGFR^+^ cells (**Fig. 2C**). To reduce bystander untransduced (UT) T cell contamination in immune assays, NGFRt^+^ cells were magnetically enriched, resulting in a ∼90% pure CAR^+^ T cell population based on frequency of Myc-tag+ and NGFR+ cells (**Fig. 2D**). MUC4 CAR T cells were used for experiments 10-12 days after initial T cell isolation and activation (**Fig. 2B**). As expected, we saw a pronounced decrease in T cell number and viability 48 hours after T cell stimulation with anti-CD3/CD28 and IL-2, the peak of T cell activation (**Fig. 2E**). Nevertheless, over the course of one week, both UT and MUC4 CAR transduced T cells expanded ∼30-fold and regained ∼90% viability (**Fig. 2F**), indicating that MUC4 CAR expression alone is not toxic to T cells. Importantly, MUC4 CAR T cells continued to expand and maintained high cell viability following NGFR magnetic selection (**Fig. 2G**), establishing our CAR T cell generation workflow.

Next, we assessed MUC4 CAR antigen-specific cytotoxicity *in vitro*. MUC4 CAR-T cells were co-cultured with MUC4-negative HEK293T cells as a negative specificity control and with MUC4-expressing CRC lines T84 (metastatic) and HT29-MTX (methotrexate-resistant) (**Fig. 3A**). Across multiple effector-to-target (E:T) ratios (0.5:1 to 4:1), HEK293T cells remained resistant to CAR-T cell-mediated killing, indicating minimal off-target activity. In contrast, MUC4 CAR-T cells induced pronounced, dose-dependent cytotoxicity against both T84 and HT29-MTX cells. This selective antitumor activity was consistently confirmed across orthogonal platforms, including LDH release (**Fig. 3B**), CCK-8 viability (**Fig. 3C**), microscopic examination (**Fig. S1A**), and flow cytometric live/dead cell detection (**Fig. S1B**). Together, these data demonstrate robust antigen-restricted cytotoxic function of MUC4 CAR-T cells against methotrexate-resistant CRC targets *in vitro*.

**Figure 3.**
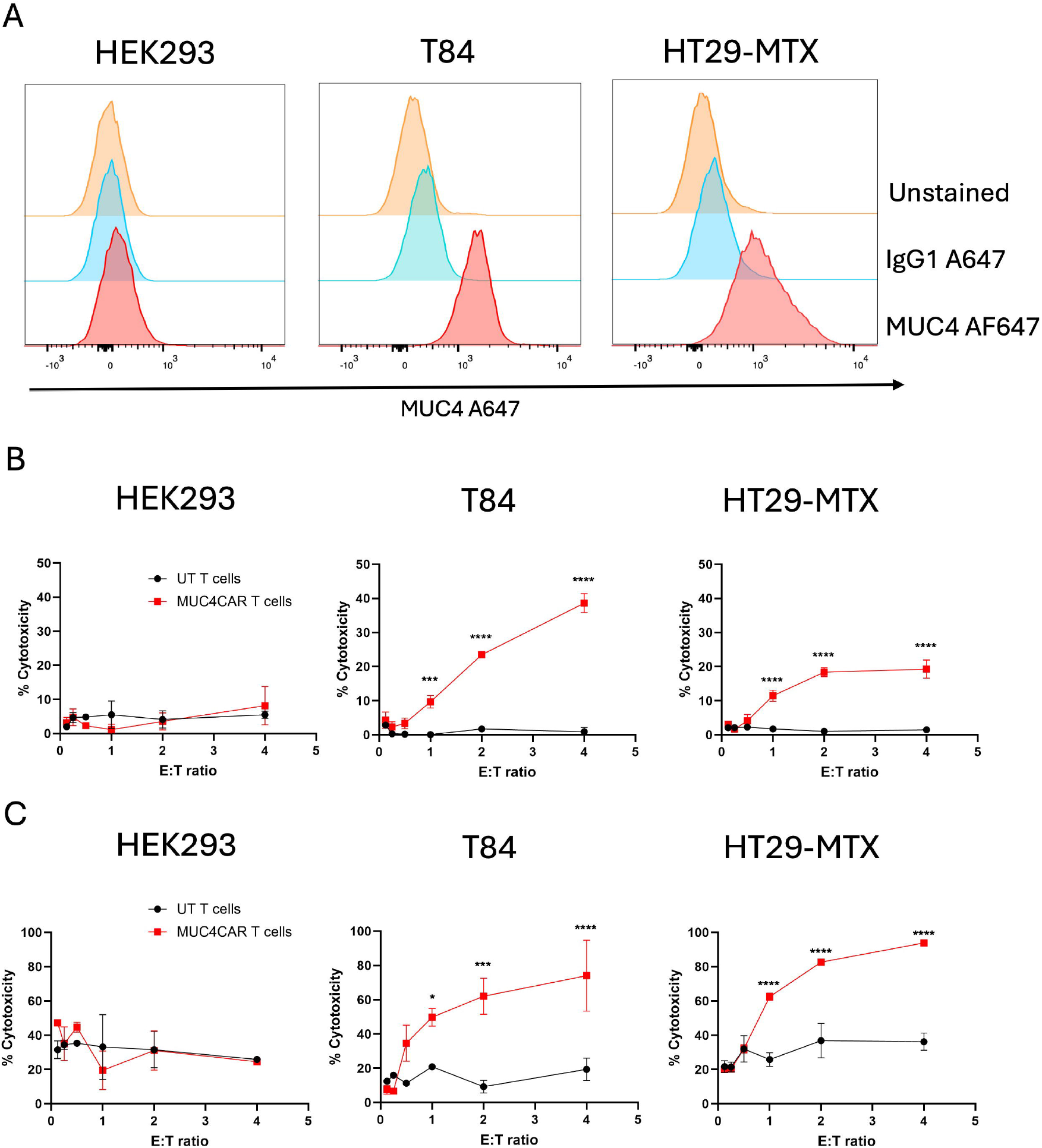
MUC4 CAR-T cells mediate potent *in vitro* cytotoxicity against HT29-MTX and T84 colorectal cancer cells. (A) Flow cytometric analysis of surface MUC4 expression in HEK293T, T84, and HT29-MTX cells. (B) LDH release assay and (C) CCK-8 viability assay showing dose-dependent cytotoxicity following co-culture of MUC4 CAR-T cells with target cells at the indicated effector-to-target (E:T) ratios. Statistical analyses were performed using two-way ANOVA. *, ***, and **** denote p < 0.05, p < 0.0001, and p < 0.00001, respectively. Data are representative of three independent experiments using three donors and are presented as mean ± SEM.

### MUC4 CAR-T cells suppress localized and metastatic methotrexate-resistant HT29-MTX tumor growth *in vivo*

Having established potent *in vitro* cytotoxic activity against MUC4^+^ chemoresistant target cells, we next evaluated the *in vivo* efficacy and safety of MUC4 CAR-T cells. We initially tested subcutaneous (s.c.) xenograft NSG mouse models using HT29-MTX and T84 cells. While T84 cells demonstrated poor and inconsistent subcutaneous engraftment, HT29-MTX tumors grew robustly and were therefore prioritized for *in vivo* efficacy studies (data not shown).

Intravenous (i.v.) adoptive transfer of MUC4 CAR-T cells, but not UT T cells, into HT29-MTX s.c. tumor-bearing NSG mice significantly attenuated tumor progression (**Fig. 4A**). Tumor growth kinetics analyzed by mixed-effects modeling (time × treatment) demonstrated a highly significant treatment effect (p < 0.0001), which was independently supported by area-under-the-curve (AUC) analysis (**Fig. 4B**). No adverse side effects caused by MUC4 CAR-T cell injection were noted. These data indicate that MUC4 CAR-T cells effectively restrain the growth of localized methotrexate-resistant CRC tumors *in vivo*.

**Figure 4.**
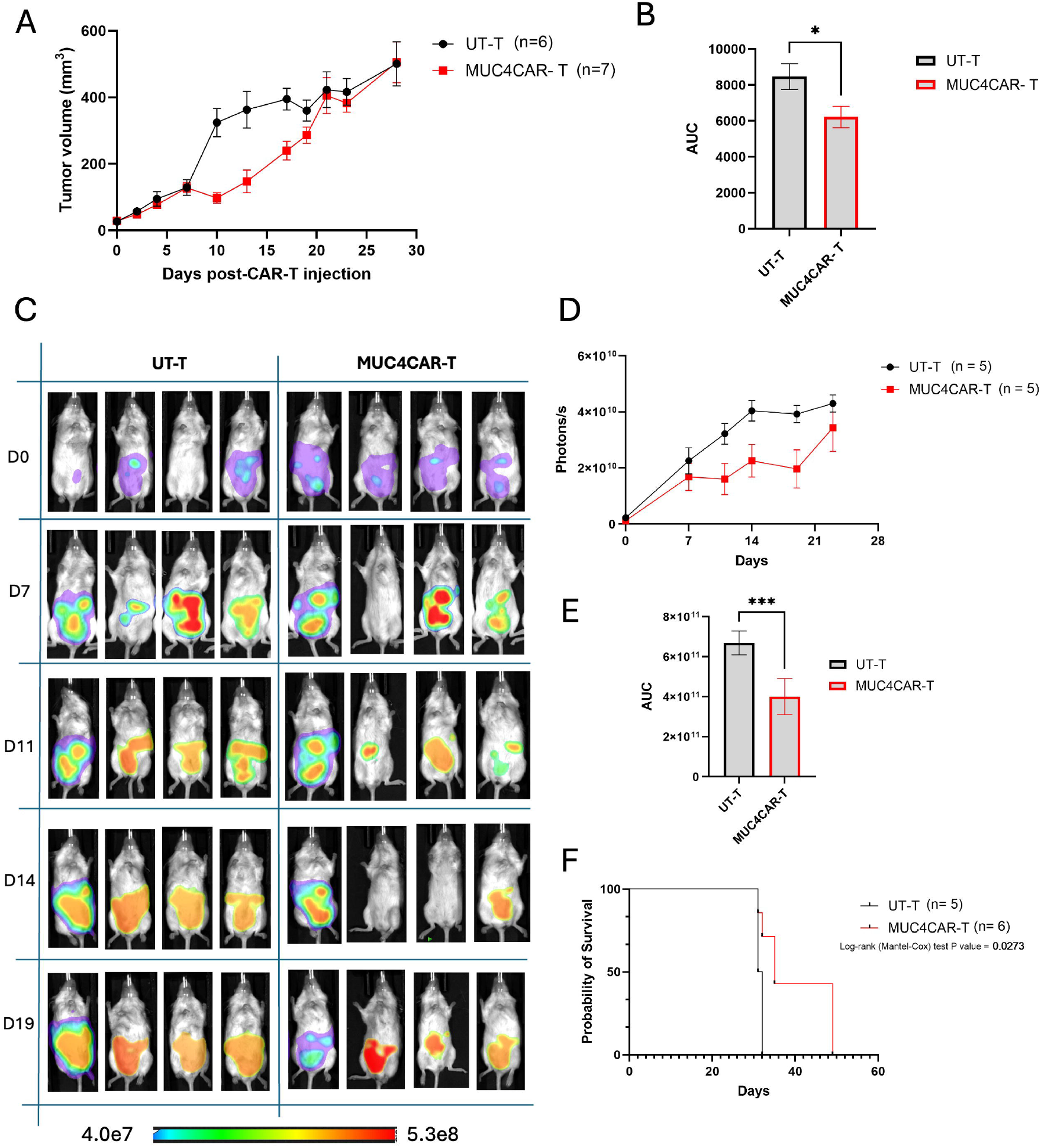
MUC4 CAR-T cells restrain chemotherapy-resistant HT29-MTX tumor growth and improve survival in NSG mice. All *in vivo* studies were performed in 8–12-week-old female NSG mice. (A) Subcutaneous xenograft model: 5 × 10□ HT29-MTX cells were implanted, followed by intravenous administration of 5 × 10□ MUC4 CAR-T cells on day 7. Tumor growth was monitored longitudinally by caliper measurements. (B) Area-under-the-curve (AUC) quantification of tumor growth from (A). (C) Disseminated xenograft model: 5 × 10^6^ HT29-MTX-Luciferase cells were injected intraperitoneally, and tumor burden was monitored by bioluminescence imaging following intraperitoneal administration of D-luciferin (150 mg/kg) using the AMI imaging system. Intravenous administration of 5 × 10^6^ MUC4 CAR-T cells on day 7. (D) Longitudinal disseminated tumor burden kinetics following MUC4 CAR-T treatment. (E) AUC quantification of the bioluminescence data in (D). (F) Kaplan–Meier survival analysis with log-rank test from the experiment in (C). Statistical analyses in (B) and (E) were performed using unpaired t-tests; * and *** denote p < 0.05 and p < 0.0001, respectively. Data are presented as mean ± SEM.

To enable longitudinal and anatomically relevant monitoring of metastatic CRC, we generated luciferase-expressing variants of T84 and HT29-MTX cell lines. Both lines were lentivirally transduced with a dual firefly luciferase–GFP reporter separated by a 2A peptide. A multiplicity of infection (MOI, number of virus particles per cell) of 10 achieved uniform GFP reporter integration without compromising viability (**Fig. S2A**,**B**). GFP^+^ cells were purified by FACS (**Fig. S2C**) and expanded, maintaining stable GFP reporter expression over serial passages (**Fig. 2D**), consistent with durable transgene integration. Functional validation confirmed robust bioluminescent signal following incubation with D-luciferin substrate (**Fig. S2E**), supporting their suitability for *in vivo* tracking. Crucially, MUC4 surface expression levels were not affected compared to the parental cell lines (**Fig. S2F**). The peritoneum is a common metastatic site for advanced CRC and is associated with poor outcomes (18). We thus sought to establish intraperitoneal CRC xenograft models to better recapitulate the metastatic behavior of mucinous CRC. Consistent with the sex of the cancer patient of origin, T84-Luc-GFP cells were implanted into male NSG mice and HT29-MTX-Luc-GFP cells into female NSG mice at two different cell doses, 2×10^6^ and 5×10^6^ cells (**Fig. S3A**). Longitudinal bioluminescence imaging revealed markedly superior and more consistent engraftment of HT29-MTX-Luc-GFP cells compared with T84-Luc-GFP cells (**Fig. S3B**,**C**), prompting selection of the HT29-MTX model for subsequent studies.

Administration of MUC4 CAR-T cells i.v. significantly delayed HT29-MTX disseminated tumor progression, as assessed by luciferase activity (**Fig. 4C,D**) and calculated using AUC analysis (**Fig. 4E**), and significantly improved the overall survival of tumor-bearing mice (**Fig. 4F**). Gross pathological analysis following mouse necropsy at 3 weeks post MUC4 CAR T cell i.v. infusion revealed no differences in body weight or organ weight normalized to body weight across saline, UT T cell, and MUC4 CAR T cell groups (**Fig. S4**). Frequent tumor engraftment in the diaphragm, pancreas, and uterus was observed in saline and UT T cell treated animals, whereas MUC4 CAR-T cell-treated mice exhibited substantially reduced tumor burden at these sites.

Collectively, these findings demonstrate that MUC4 CAR-T cells exert robust antitumor activity against localized methotrexate-resistant CRC *in vivo*, retain efficacy in a clinically relevant disseminated methotrexate-resistant CRC model, and are safe.

## Discussion

CAR-T cell therapy is being actively pursued in CRC as a strategy to overcome chemotherapy resistance through antigen-specific tumor cell elimination. Several CAR targets have been explored in CRC, including GUCY2C, CEA, EpCAM, HER2, EGFR, and CLDN18.2 (19). However, as is observed across solid tumors in general (9,10), CAR-T cell strategies targeting conventional epithelial antigens in CRC have shown limited clinical success due to antigen heterogeneity, on-target/off-tumor toxicity arising from shared expression in normal tissues, and poor persistence within the immunosuppressive tumor microenvironment. In addition, specific physical barriers of mucin-rich CRC tumors impair immune cell infiltration, effective synapse formation, and lead to immunotherapy failures (3). These limitations highlight the need for alternative strategies that are both tumor-restricted and biologically relevant to disease progression.

Mucins are high-molecular-weight, heavily glycosylated proteins forming protective, lubricating, and signaling barriers on epithelial surfaces, categorized into two main types: secreted and cell-surface associated. Secreted (gel-forming) mucins, such as MUC2 and MUC5AC, form a mucus layer, while membrane-bound mucins like MUC1, MUC4, and MUC16 anchor to the cell apical surface, playing roles in pathogen binding and cell signaling (20). Membrane-bound mucin-directed immunotherapy in solid tumors has primarily focused on MUC1 and, more recently, MUC16; however, each presents translational constraints that have tempered clinical success (21,22). MUC1 is overexpressed and depolarized in many carcinomas but also has appreciable expression in a range of normal epithelia and hematopoietic cells, whereas MUC16 is normally expressed on the ocular surface as protection from pathogens (23,24). Furthermore, MUC16 is often proteolytically cleaved at the extracellular domain and shed from the cell surface, which limits durable CAR engagement and anti-tumor efficacy (21,25). In contrast, MUC4 displays several attributes that support its candidacy as a tumor-associated antigen in mucinous malignancies. MUC4 expression is frequently upregulated and depolarized in aggressive epithelial tumors, including subsets of colorectal cancer (CRC), while remaining comparatively restricted in many normal tissues. Functionally, MUC4 contributes to tumor progression, metastatic fitness, chemotherapy-resistance, and immune evasion through steric shielding and pro-survival signaling (3). These features position MUC4 not only as a surface marker but also as a biologically relevant vulnerability against aggressive tumors.

In this study, MUC4 CAR-T cells significantly delayed tumor progression in both subcutaneous and disseminated models of methotrexate-resistant CRC. Of note, we established an *in vivo* model of human CRC peritoneal invasion. Peritoneal cancer invasion presents significant challenges due to late-stage diagnosis, rapid spread throughout the abdominal cavity, and high recurrence rates, greatly limiting patients’ current treatment options (18). The ability to restrain the growth of mucinous, standard-of-care therapy-refractory tumors highlights the potential of this approach to address a critical therapeutic gap where conventional modalities often fail.

However, HT29-MTX CRC tumors were controlled rather than fully eradicated by MUC4 CAR-T cells, indicating that additional improvements to our strategy are needed for maximal efficacy *in vivo*. Multiple engineering strategies may enhance the therapeutic ceiling of MUC4-directed CAR-T cells. Recent studies revealed that 4-1BB-CD3zeta CAR signaling elicits a more durable anti-tumor response than CD28-CD3zeta CAR against solid tumors (26). Therefore, a future comparative evaluation of CD28-CD3zeta (used in the present study) vs. 4-1BB-CD3zeta domains in the MUC4 CAR design could clarify whether metabolic fitness, persistence, and ultimately efficacy can be improved through alternative signaling programs. In parallel, transcriptional reprogramming approaches such as FOXP3 upregulation, previously implicated in promoting resilience within suppressive microenvironments, may augment functional stability and tumor microenvironment fitness of CAR-T cells in mucin-rich tumors (27). Our findings support further exploration of CAR-T cell persistence-enhancing designs in the setting of CRC.

Importantly, MUC4 overexpression extends beyond colorectal cancer to other aggressive epithelial malignancies. Most prominent examples are pancreatic ductal adenocarcinoma (28), which similarly to CRC exhibits peritoneal and diaphragmatic dissemination (29), and ovarian cancer, with MUC4 expression being significantly higher than either MUC1 or MUC16 expression in early-stage ovarian tumors with 100% incidence and in in advanced stage ovarian tumors (30). These features broaden the potential translational scope of MUC4-targeted CAR-T therapy across multiple mucin-driven tumors.

Altogether, our findings provide preclinical support that MUC4-directed CAR-T cells can recognize and eliminate chemotherapy-resistant CRC targets, supporting the prioritization of MUC4 as a therapeutic target in mucin-rich solid tumors.

## Acknowledgments

This work was supported by Swim Across America Grant 23-1579 to LMRF and an MUSC Specialized Center of Research Excellence (SCORE) 5U54DA016511-18 Pilot Project Award to AC. This study was supported in part by the Flow Cytometry and Cell Sorting Shared Resource, Hollings Cancer Center, Medical University of South Carolina (P30 CA138313).

