## Supplement for "MUC4-targeted CAR-T Cells Restrain Chemotherapy-resistant Colorectal Cancer"

Fig. S1

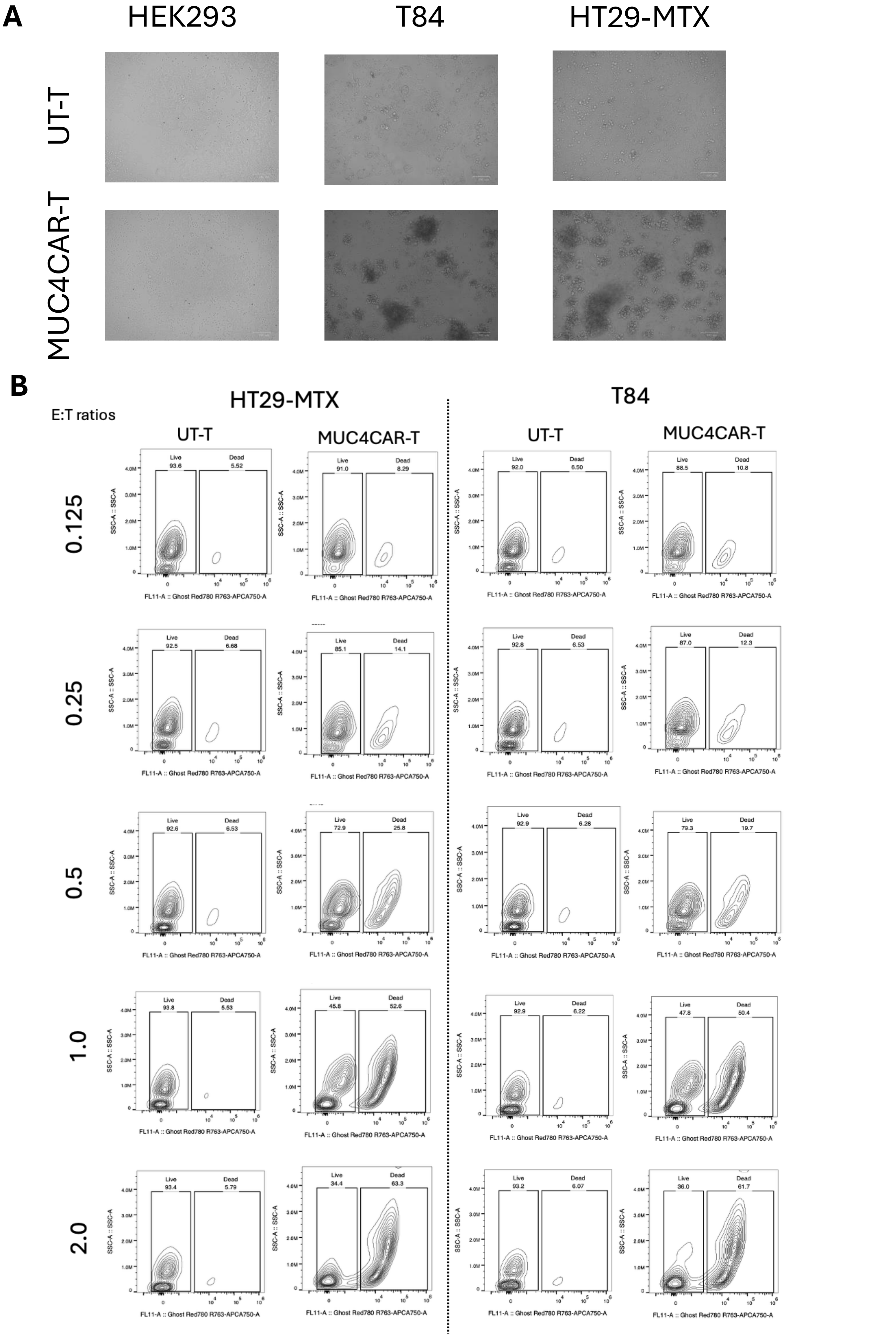

A
HEK293
T84
HT29-MTX
UT-T
MUC4CAR-T
B
HT29-MTX
T84
E:T ratios
UT-T
MUC4CAR-T
UT-T
MUC4CAR-T
0.125
Live 93.6 Dead 5.52
Live 91.0 Dead 8.29
Live 92.0 Dead 6.50
Live 88.5 Dead 10.8
0.25
Live 92.5 Dead 6.68
Live 85.1 Dead 14.1
Live 92.8 Dead 6.53
Live 87.0 Dead 12.3
0.5
Live 92.6 Dead 6.53
Live 72.9 Dead 25.8
Live 92.9 Dead 6.28
Live 79.3 Dead 19.7
1.0
Live 93.8 Dead 5.53
Live 45.8 Dead 52.6
Live 92.9 Dead 6.22
Live 47.8 Dead 50.4
2.0
Live 93.4 Dead 5.79
Live 34.4 Dead 63.3
Live 93.2 Dead 6.07
Live 36.0 Dead 61.7
SSC-A :: SSC-A
FL11-A :: Ghost Red780 R763-APCA750-A

**Supplementary Figure 1. Microscopy and flow cytometry confirm MUC4 CAR-T–mediated killing of HT29-MTX and T84 cells.** (A) Representative brightfield images of tumor cells co-cultured with MUC4 CAR-T cells at an effector-to-target (E:T) ratio of 4:1 for 36 hours. (B) Flow cytometric quantification of Ghost Red 780–positive dead cells across the indicated effector-to-target (E:T) ratios.

Fig. S2

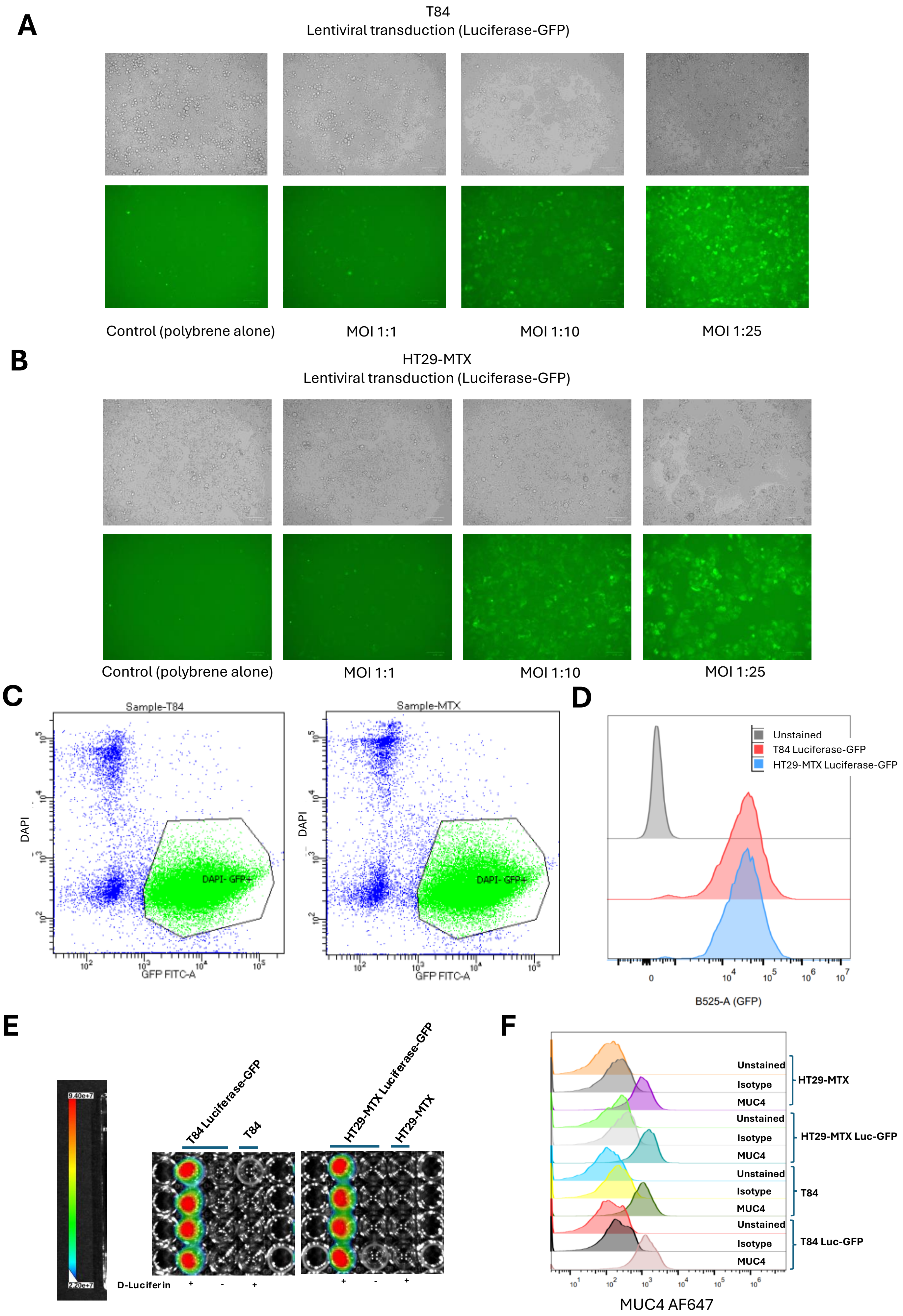

A
T84
Lentiviral transduction (Luciferase-GFP)
Control (polybrene alone)
MOI 1:1
MOI 1:10
MOI 1:25
B
HT29-MTX
Lentiviral transduction (Luciferase-GFP)
Control (polybrene alone)
MOI 1:1
MOI 1:10
MOI 1:25
C
Sample-T84
Sample-MTX
DAPI
DAPI- GFP+
GFP FITC-A
D
Unstained
T84 Luciferase-GFP
HT29-MTX Luciferase-GFP
B525-A (GFP)
E
T84 Luciferase-GFP
T84
HT29-MTX Luciferase-GFP
HT29-MTX
D-Luciferin
F
Unstained
Isotype
MUC4
HT29-MTX
HT29-MTX Luc-GFP
T84
T84 Luc-GFP
MUC4 AF647

**Supplementary Figure 2. Stable transduction and characterization of luciferase-GFP-expressing colorectal cancer cell lines.** HT29-MTX and T84 cells were lentivirally transduced with plasmid pLV-EF1A:Luciferase:T2A:EGFP at a titer of $2.93 \times 10^8$ viral particles/mL using multiplicities of infection (MOI) of 1, 10, and 25 in the presence of polybrene in DMEM + 10% FBS. (A, B) Representative bright-field and fluorescence images of GFP expression in HT29-MTX (A) and T84 (B) cells at 72 h post-transduction. (C) $GFP^+$ $DAPI^-$ cells were sorted using FACSAria III™ cell sorter one week after transduction. (D) Retention of GFP expression was evaluated after four passages post-sorting by flow cytometry (Beckman Coulter CytoFLEX™). (E) Luciferase activity was confirmed by bioluminescence imaging following the addition of D-luciferin (1:100 dilution from a 15 mg/mL stock) and 5 min incubation in DMEM + 10% FBS. (F) Surface expression of MUC4 was validated after seven passages post-sorting by flow cytometry using anti-human MUC4 antibody (clone 8G7) and goat anti-mouse Alexa Fluor 647 secondary antibody.

Fig. S3

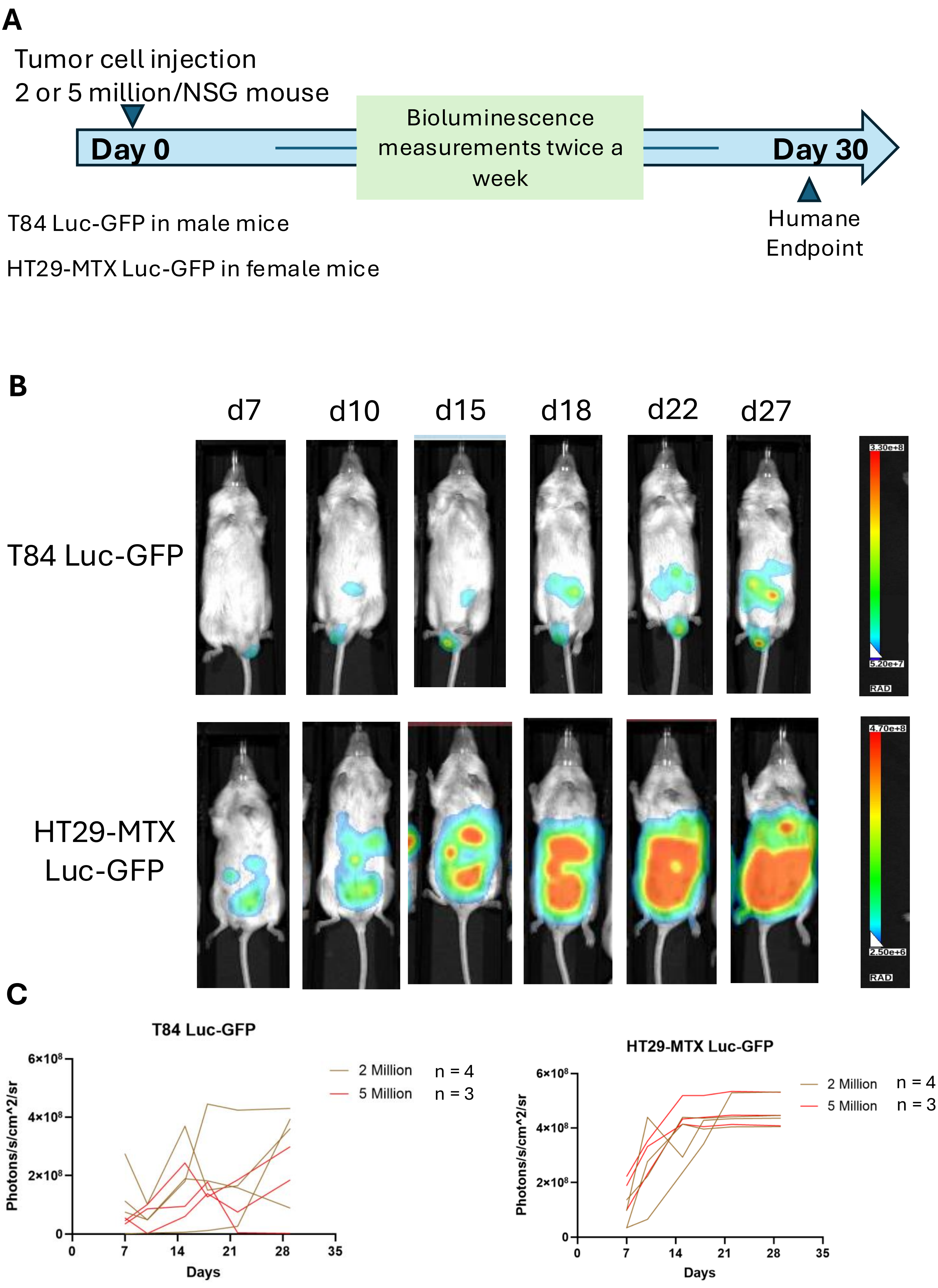

A
Tumor cell injection
2 or 5 million/NSG mouse
Day 0
Bioluminescence measurements twice a week
Day 30
Humane Endpoint
T84 Luc-GFP in male mice
HT29-MTX Luc-GFP in female mice
B
d7
d10
d15
d18
d22
d27
T84 Luc-GFP
HT29-MTX Luc-GFP
3.30e+8
5.20e+7
RAD
4.70e+8
2.50e+6
RAD
C
T84 Luc-GFP
Photons/s/cm^2/sr
Days
2 Million n = 4
5 Million n = 3
HT29-MTX Luc-GFP
Photons/s/cm^2/sr
Days
2 Million n = 4
5 Million n = 3

**Supplementary Figure 3. *In vivo* engraftment and monitoring of luciferase-expressing colorectal tumor cells in NSG mice.** (A) Schematic workflow outlining tumor engraftment and subsequent readout measurements. (B) Longitudinal bioluminescence imaging of tumor growth for intraperitoneal HT29-MTX-luc-GFP and T84-luc-GFP. (C) Quantification of tumor burden progression for two cell doses ($2 \times 10^6$ and $5 \times 10^6$ cells) of two cell lines (HT29-MTX-luc-GFP and T84-luc-GFP) represented as maximum radiance values over time. N represents the number of mice.

Fig. S4

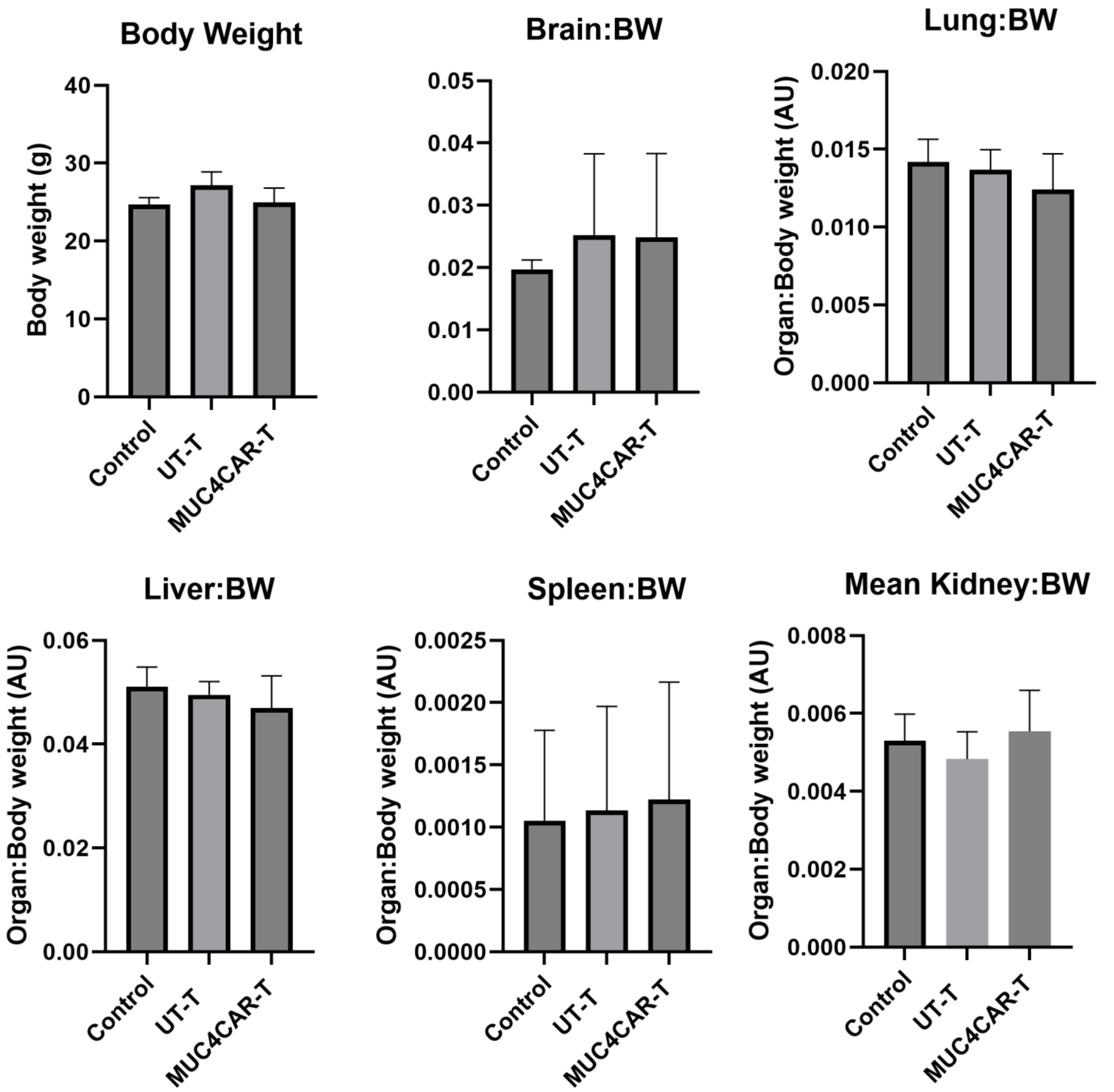
Body Weight
Body weight (g)
40
30
20
10
0
Control
UT-T
MUC4CAR-T
Brain:BW
0.05
0.04
0.03
0.02
0.01
0.00
Control
UT-T
MUC4CAR-T
Lung:BW
Organ:Body weight (AU)
0.020
0.015
0.010
0.005
0.000
Control
UT-T
MUC4CAR-T
Liver:BW
Organ:Body weight (AU)
0.06
0.04
0.02
0.00
Control
UT-T
MUC4CAR-T
Spleen:BW
Organ:Body weight (AU)
0.0025
0.0020
0.0015
0.0010
0.0005
0.0000
Control
UT-T
MUC4CAR-T
Mean Kidney:BW
Organ:Body weight (AU)
0.008
0.006
0.004
0.002
0.000
Control
UT-T
MUC4CAR-T

**Supplementary Figure 4. MUC4 CAR-T cell treatment does not impact body or organ weight in peritoneally disseminated HT29-MTX-luc-GFP bearing NSG mice.** Total body weight (BW), as well as weight normalized to BW for brain, lung, liver, spleen, and kidneys across control (saline), UT T cell, and MUC4 CAR-T cell treated mice.
